# Yeast surface display of full-length human microtubule-associated protein tau

**DOI:** 10.1101/624429

**Authors:** Shiyao Wang, Yong Ku Cho

**Affiliations:** Department of Chemical and Biomolecular Engineering; Department of Biomedical Engineering, Institute for Systems Genomics, CT Institute for the Brain and Cognitive Sciences, University of Connecticut, Storrs, CT 06269, United States

**Keywords:** microtubule-associated protein tau, *Saccharomyces cerevisiae*, yeast surface display, antibody, conformation

## Abstract

Microtubule-associated protein tau is an intrinsically-disordered, highly soluble protein found primarily in neurons. Under normal conditions, tau regulates the stability of axonal microtubules and intracellular vesicle transport. However, in patients of neurodegeneration such as Alzheimer’s disease (AD), tau forms neurofibrillary deposits, which correlates well with the disease progression. Identifying molecular signatures in tau, such as post-translational modification, truncation, and conformational change has great potential to detect earliest signs of neurodegeneration, and develop therapeutic strategies. Here we show that full-length human tau, including the longest isoform found in the adult brain can be robustly displayed on the surface of yeast *Saccharomyces cerevisiae*. Yeast-displayed tau binds to anti-tau antibodies that cover epitopes ranging from the N-terminus to the 4R repeat region. Unlike tau expressed in the yeast cytosol, surface-displayed tau was not phosphorylated at sites found in AD patients (probed by antibodies AT8, AT270, AT180, PHF-1). However, yeast-displayed tau showed clear binding to paired helical filament (PHF) tau conformation-specific antibodies Alz-50, MC-1, and Tau-2. Although the tau possessed a conformation found in PHFs, oligomerization or aggregation into larger filaments were undetected. Taken together, yeast-displayed tau enables robust measurement of protein interactions, and is of particular interest for characterizing conformational change.

## Introduction

Microtubule-associated protein tau (MAPT) is highly expressed in neurons to stabilize microtubules, and is critical for regulating axonal transport and maintaining morphology (Roy et al., 2005). Direct linkage of tau to neurodegeneration was established by the identification of *MAPT* mutations that cause inherited frontotemporal dementia and Parkinsonism linked to chromosome 17 (FTDP-17) (Hutton et al., 1998; Ingram & Spillantini, 2002). Severity of dementia correlates with the level of paired helical filament (PHF) composed of tau aggregates, but not to the level of senile plaque, composed of amyloid β (Arriagada et al., 1992; Giannakopoulos et al., 2003). Site-specific post-translational modifications (PTMs), such as phosphorylation regulates the tau-microtubule interaction (Gendron & Petrucelli, 2009; Geschwind, 2003) and axonal transport (Ebneth et al., 1998; Ishihara et al., 1999; Tatebayashi et al., 2004). Various tau phosphorylation sites specific to patients with neurodegeneration, including Alzheimer’s disease, Pick’s disease, and progressive supranuclear palsy have been discovered (Ludolph et al., 2009; Morris et al., 2015; Spires-Jones et al., 2009). On the other hand, negatively charged surfaces and polymers also drives conformational change in tau in the absence of phosphorylation (Friedhoff et al., 1998; Goedert et al., 1996; Hasegawa et al., 1997; Kampers et al., 1996; Wilson & Binder, 1997).

An essential tool for detecting tau is anti-tau antibodies. According to the online database by Alzforum (alzforum.org), >500 anti-tau antibodies are currently available. Tau is highly soluble and intrinsically disordered, allowing generation of anti-tau antibodies using peptide antigens. In particular, peptide antigens are widely used to generate antibodies that target site-specific phosphorylation in tau. Existing methods for characterizing anti-tau antibodies also primarily rely on synthesized peptides (Ercan et al., 2017; Mercken et al., 1992; Petry et al., 2014). However, approaches using peptides cannot capture the conformation sensitivity of anti-tau antibodies, which in many cases depend on recognizing multiple discontinuous epitopes in tau (Carmel et al., 1996; Jicha, Bowser, et al., 1997). It has also been demonstrated that phospho-tau antibodies can be specific to conformation found in PHFs (Jicha, Lane, et al., 1997). To fully probe the conformation sensitivity and specificity of anti-tau antibodies, an approach that provides full-length tau with high-throughput genotype-phenotype linkage would be ideal.

Here we show that full-length human tau can be robustly displayed on the surface of yeast *Saccharomyces cerevisiae*. We find that unlike human tau expressed in the yeast cytosol (Rosseels et al., 2015; Vandebroek et al., 2005), yeast surface displayed tau is primarily non-phosphorylated, and allow direct assessment of antibody binding. Many linear epitopes in tau, ranging from the N-terminus to the 4R repeat region were available for antibody binding. Importantly, yeast-displayed tau showed conformation recognized by antibodies specific to PHF conformation, potentially due to the negatively charged environment of the yeast surface. Moreover, although the tau possessed a conformation found in PHFs, oligomerization or aggregation into larger filaments were undetected. Therefore, yeast surface displayed tau provides a novel platform for assessing antibody binding, and also for characterizing its conformational change.

## Materials and Methods

### Antibodies

Anti-tau antibodies used in this study and the dilutions used are listed in **Table 1**.

**Table 1.** Anti-tau antibodies used for yeast cell labeling.

| **Antibody** | **Epitope** | **Supplier (catalog#)** | **Dilution –**<br>**yeast cell**<br>**labeling** |
| --- | --- | --- | --- |
| **1E1/A6** | 275-291 | Millipore<br>(05-804) | 1:100 |
| **Alz50** | Conformation-specific | Peter Davies | 1:50 |
| **AT180** | Thr231 | Thermo Fisher<br>(MN1040) | 1:100 |
| **AT270** | Thr181 | Thermo Fisher<br>(MN1050) | 1:100 |
| **AT8** | Ser202-Thr205 | Thermo Fisher<br>(MN1020) | 1:100 |
| **HT7** | 159-163 | Thermo Fisher<br>(MN1000) | 1:100 |
| **MC-1** | Conformation-specific | Peter Davies | 1:50 |
| **PHF-1** | Ser396 Ser404 | Peter Davies | 1:50 |
| **PHF-6** | Thr231 | Thermo Fisher<br>(35-5200) | 1:100 |
| **T22** | Oligomer-specific | Millipore<br>(ABN454-I) | 1:100 |
| **Tau-1** | Non-phosphorylated<br>Ser195/198/199/202 | Millipore<br>(MAB3420) | 1:100 |
| **Tau-2** | Conformation-specific | Sigma<br>(T5530) | 1:100 |
| **Tau-5** | 210-230 | Thermo Fisher<br>(MA5-12808) | 1:100 |
| **Tau-46** | 428-441 | Sigma<br>(T9450) | 1:100 |
| **TOMA-1** | Oligomer-specific | Millipore<br>(MABN819) | 1:100 |

### Cloning and yeast surface display of full-length human MAPT

DNA sequence encoding the 383 amino acid isoform (0N4R) of human microtubule-associated protein tau (*MAPT*, Gene ID: 4137) was codon optimized for yeast *S. cerevisiae* using an online tool (GeneArt), and synthesized (IDTDNA). Yeast codon-optimized DNA sequence encoding the 2 N-terminal inserts were also synthesized (IDTDNA), and inserted into the 0N4R sequence using overlap extension PCR (Zhao et al., 1998) (see **Table 2** for primers used). For yeast surface display, the DNA was cloned into the yeast surface display vector pCT-GFPM (a gift from Dr. Eric Shusta) (Pavoor et al., 2009) between restriction sites NheI and BamHI. The result construct pCT-MAPT 2N4R were used to transform *S. cerevisiae* strain EBY100 using frozen-EZ yeast Transformation II kit (Zymo Reseach, Cat. NO. T2001) and grown on SD-CAA agar plates for 3 days. Single colony of cells were picked up and grown in 3 mL of SD-CAA media at 30 °C, 250 rpm for overnight, pelleted, and re-suspended in 3 mL of SG-CAA at 20 °C, 250 rpm for 20 hours to induce the expression of tau.

**Table 2.**
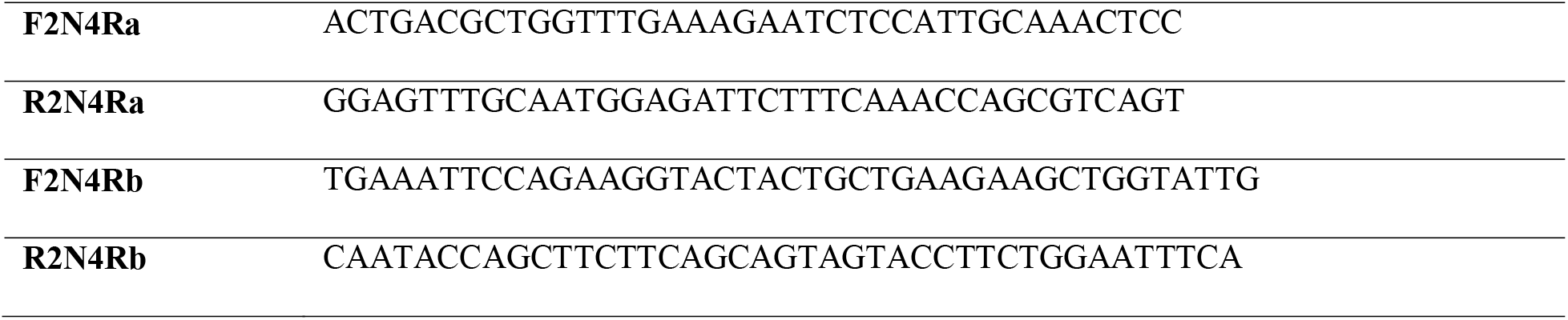
Primers for cloning pCT-MAPT-2N4R

### Deglycosylation of yeast-displayed tau

Deglycosylation of yeast surface displayed tau was performed using PNGase (New England Biolabs (NEB), Cat. No. P0704S) and α1-2,3,6 Mannosidase (NEB, Cat. No. P0768S). 0.5 × 10^7^ of yeast cells were washed, pelleted, and resuspended in 50 μL of phosphate buffered saline (PBS: 8 g/L NaCl, 0.2 g/L KCl, 1.44 g/L Na_2_HPO_4_, 0.24 g/L KH_2_PO_4_) pH 7.4 with 0.1 % bovine serum albumin (BSA: 1 g/L), 1 % NP-40, 5 μL of 10 × GlycoBuffer 2 (supplied by NEB, 500 mM Sodium Phosphate, pH 7.5), and 1 μL of PNGase. After incubation for 2 hours at 30 °C, cells were washed 3 times with PBS pH 7.4 with 0.1 % BSA, and then incubated 2 hours at 30 °C in 50 μL of PBS pH 7.4 with 0.1 % BSA, 1 × Zinc (supplied by NEB), 5 μL of 10 × GlycoBuffer 4 (supplied by NEB, 500 mM Sodium Phosphate, pH 4.5), and 1 μL of α1-2,3,6 Mannosidase.

### Antibody labeling of yeast-displayed tau

0.5 × 10 of yeast cells were washed, pelleted, and resuspended in 100 μL of PBS pH 7.4 with 0.1 % BSA and the primary antibody at the recommended dilution rate (listed in **Table 1**). After the primary incubation for one hour on ice, cells were washed three time with 500 μL PBS pH 7.4 with 0.1% BSA (5 mins incubation on ice each time). Then, cells were resuspended in 100 μL of PBS pH 7.4 with 0.1 % BSA and one of the following secondary antibodies: anti-mouse Alexa Fluor IgG 488 (Thermo Fisher Scientific, Cat. No. A11001, 1:500 dilution), anti-chicken IgY Alexa Fluor 488 (Thermo Fisher Scientific, Cat. No. A11039, 1:500 dilution), anti-rabbit IgG Alexa Fluor 594 (Thermo Fisher Scientific, Cat. No. A21442, 1:500 dilution), anti-mouse IgG Alexa Fluor 594 (Thermo Fisher Scientific, Cat. No. A11005, 1:500 dilution), or anti-mouse IgM Alexa Fluor 594 (Thermo Fisher Scientific, Cat. No. A21044, 1:500 dilution), and incubated on ice for 30 mins. Labeling results were detected and collected by BD Biosciences LSR Fortessa X-20 Cell Analyzer (UConn Center for Open Research Resources and Equipment).

### 3C protease cleavage of yeast-displayed tau

In order to cleave tau proteins from the yeast surface, a 3C protease cleavage site was inserted between Aga2p and tau by inserting the *MAPT* 2N4R gene into the yeast surface display vector pCT-3C BN_hIL2 (a gift from Dr. Jamie B. Spangler) (Silva et al., 2019) between restriction sites BamHI and NotI (see **Table 3** for primers used). The resulting construct pCT-3C-MAPT 2N4R were then used for displaying tau with a 3C protease site. For the 3C protease cleavage reaction, 0.5 × 10^7^ of yeast cells were washed, pelleted, and resuspended in 50 μL of reaction buffer containing: 25 mM HEPES pH 7.5, 150 mM NaCl, 1 mM EDTA, 1 mM DTT and 1 μL HRV-3C protease (Sigma, Cat. No. SAE0045-1 MG). After 16 hours incubation on a rotor under 4 °C, cells were pelleted, and only the supernatant were collected for western blot.

**Table 3.**
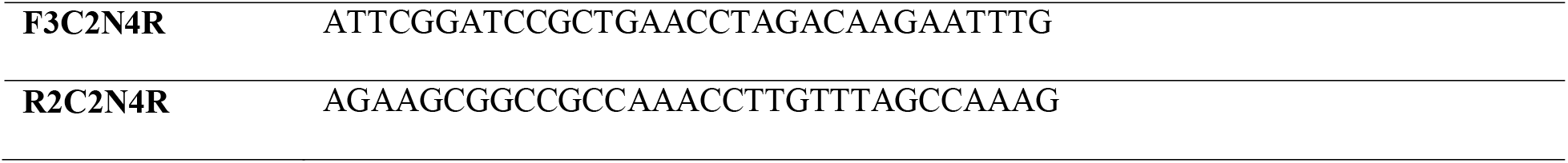
Primers for cloning pCT-3C-MAPT-2N4R

### SDS-PAGE and western blot

SDS-PAGE gels consisted of an 8 % Tris-Glycine separation gel: 4.3 mL ddH_2_O, 3.75 mL 1 M Tris pH 8.8, 2.99 ml 30 % Acrylamide/Bis 29:1 (Bio-Rad, Cat. No. 161-0156), 112.5 μL 10 % (w/v) SDS, 15 μL TEMED (Bio-Rad, Cat. No. 161-0800), and 37.5 μL 10 % (w/v) Ammonium Persulfate (APS), and a 4 % stacking gel: 2.6 mL ddH_2_O, 0.48 mL 1 M Tris pH 6.8, 0.625 ml 30 % Acrylamide/Bis 29:1, 37.5 μL 10 % (w/v) SDS, 5 μL TEMED, and 12.5 μL 10 % (w/v) APS, added in order. The protein samples were prepared with: 14 μL supernatant from the 3C protease cleavage reaction, 4 μL 5 × dissociation buffer (10 % (w/v) SDS, 6 % (w/v) DTT in 400 mM Tris pH 6.8), and 2 μL 10 × sample dye (0.5 % (w/v) Bromophenol blue in 80 % (v/v) Glycerol). Gels were then transferred to nitrocellulose membrane (Sigma, Cat. No.GE10600006), and the membranes were blocked overnight with 5 % (w/v) milk in TBST (20 mM Tris pH 7.6, 137 mM NaCl, and 1 % (v/v) Tween 20). Western blot was conducted using the Tau-13 antibody (dilution rate 1:2,000) followed by an anti-mouse IgG antibody conjugated to HRP (Thermo Fisher Scientific, Cat. No. A16066, dilution rate 1:2,000). Signals were detected using the Amersham™ hyperfilm ECL™ (Sigma, Cat. No. GE28-9068-35) and Amersham™ ECL™ western blotting reagent (Sigma, Cat. No. RPN2109).

### Thioflavin S staining

1 % (w/v) thioflavin S solution were prepared in distilled water and filted before use. 0.5 × 10^7^ of yeast cells were incubated in 500 μL thioflavin S solution for 10 mins with tubes covered by aluminum foil and then washed 3 times with 75 % (v/v) ethanol. After the incubation and washing, cells were protected from light until fluorescence signals were detected with BD Biosciences LSRFortessa X-20 Cell Analyzer using UV excitation (350 nm) and green fluorescence emission (530 nm), to detect amyloid aggregates (Espargaro et al., 2012).

### Statistical analysis

All data points were plotted whenever possible. Statistical analysis was conducted using the GraphPad Prism 8 software. When analysis of variance (ANOVA) was conducted, *post hoc* analysis was conducted to assess statistical significance. Exact test used for each dataset is specified in the figure legend.

## Results and Discussion

### Yeast surface display of full-length human tau

Human microtubule-associated protein tau (MAPT) exists in six distinct isoforms in adult tissue, generated by alternative splicing of N-terminal inserts (N1 and N2) and repeats (R1-R4) (Goedert et al., 1989). For yeast surface display, we cloned two isoforms: isoform 2N4R (441 amino acids, longest isoform among those found in adult tissue) contains both N-terminal inserts and all of repeats, while isoform 0N4R (383 amino acids), is missing the two N-terminal inserts to the C-terminus of Aga2p (**Fig. 1a**). When we expressed the 2N4R fusion construct (**Fig. 1b**) in the *S. cerevisiae* strain EBY100, antibody staining to the c-myc epitope tag inserted at the C-terminus of tau was clearly detected (**Fig. 1c**). A human tau-specific antibody recognizing all tau isoforms (Tau-5) also clearly labeled these cells (**Fig. 1c**). These results indicate expression of full-length human tau on the yeast cell surface. To quantify the expression of human tau, Tau-5 antibody binding to yeast cells expressing tau isoforms 2N4R, 0N4R, or a control protein (green fluorescent protein) was analyzed using flow cytometry. Strong binding specific to tau displaying yeast was observed (**Fig. 1d**), indicating that yeast cells can display either isoforms of human tau at similar levels.

**Figure 1.**
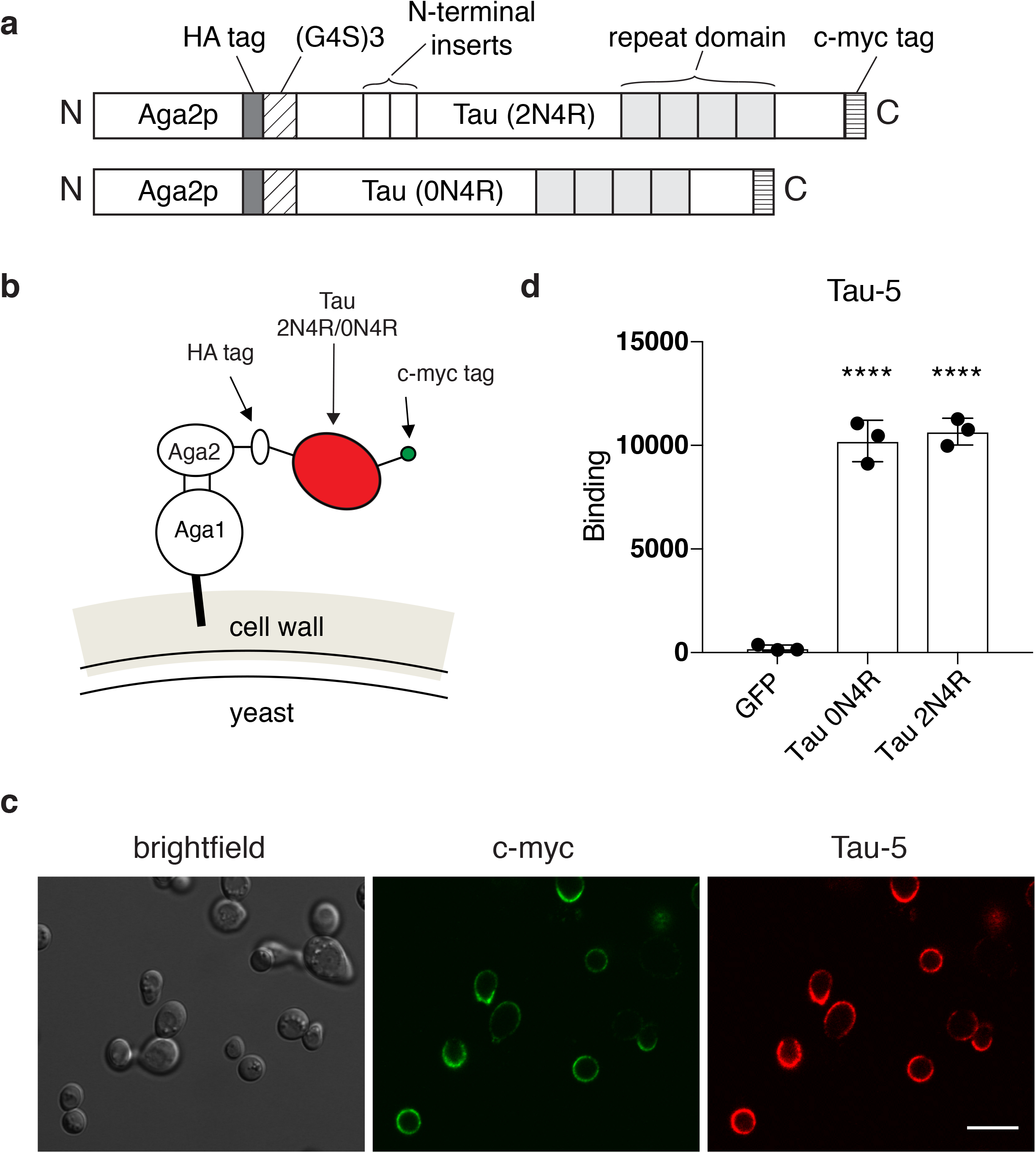
Yeast surface display of full-length human tau. (**a**) Schematics of tau display constructs generated. Human tau (2N4R or 0N4R isoform) was fused to the C-terminus of the yeast cell wall protein Aga2p. A (G_4_S)_3_ linker was inserted between Aga2p and tau. A hemagglutinin (HA) epitope tag and a c-myc epitope tag were inserted to the N- and C-terminus of tau, respectively. (**b**) A schematic representation of yeast surface displayed tau. (**c**) Confocal microscopy images of yeast surface displayed 2N4R tau. C-myc tag (green) was probed using a chicken anti-c-myc antibody, followed by an anti-chicken IgY conjugated with Alexa 488. Human tau (red) was probed using the anti-human tau antibody Tau-5, followed by an antimouse IgG conjugated with Alexa 596. Scale bar, 10 μm. (**d**) Flow cytometry quantification of fluorescence intensity of Tau-5 binding to yeast cells displaying GFP, tau 0N4R, and tau 2N4R. Data points are geometric means and error bars indicate standard deviation from three independent culture preparations. Statistics for panel (**d**): One-way ANOVA followed by Dunnett’s multiple comparisons test, comparing Tau-5 binding in GFP vs. tau 0N4R or tau 2N4R cells, **** *P* < 0.0001.

### Glycosylation of yeast-displayed tau

Heterologous proteins expressed in *S. cerevisiae* are often glycosylated. In particular, high level of *N*-glycosylation has been consistently observed in recombinant proteins expressed in the yeast *S. cerevisiae* (Parthasarathy et al., 2006; Smith et al., 1985; Tang et al., 2016). *N*-glycosylation occurs in asparagine residues, and a sequence based prediction (Chauhan et al., 2013) identified three potential N-glycosylation sites, N166, N358, and N409 in 2N4R tau (residue scores > 0.8). The same prediction approach also identified 4 potential *O*-glycosylation sites, T75, T174, S209, and T211 in 2N4R tau (residue scores > 0.4). To assess the glycosylation status of yeast-displayed tau, we generated a construct with a human rhinovirus 3C protease recognition site inserted at the N-terminus of tau (**Fig. 2a**). After yeast display, we treated the cells with a recombinant 3C protease, separated the cleaved proteins using SDS-PAGE, and performed western blot using the anti-tau antibody Tau-13 (**Fig. 2b**). Since the buffer used for 3C protease cleavage contains dithiothreitol (DTT), the buffer by itself would result in reduction of the disulfide bond between Aga1p and Aga2p, releasing the Aga2p-tau fusion protein. When we incubated 2N4R tau displaying yeast cells with the 3C protease buffer, a large molecular weight smear was detected in tau displaying cells (**Fig. 2b**, lane 2), while nothing was detected in cells displaying a control protein (antibody single-chain variable region fragment, **Fig. 2b**, lane 1). When we treated tau displaying cells with 3C protease without deglycosylation, cleavage products were detected between molecular weights of 55 to 70 kDa (**Fig. 2b**, lane 3). In tau displaying cells treated with PNGase, which cleaves most *N*-glycosylation, and α-mannosidase for removing parts of *O*-glycosylation, substantial molecular weight shift was observed (**Fig. 2b**, lane 4), indicating deglycosylation. In this product, a major band appeared at 58 kDa with additional bands (**Fig. 2b**, lane 4). We also ran a mixture of 6 tau isoforms expressed from *E. coli* (**Fig. 2b**, lane 6), the largest of which corresponds to the 2N4R tau. The major deglycosylated, 3C protease cleavage product (58 kDa) was substantially smaller than the largest isoform expressed from *E. coli* (67 kDa). Even though the human rhinovirus 3C protease is relatively sequence specific (Ullah et al., 2016), full-length tau may be highly susceptible for erroneous cleavage due to its intrinsically disordered nature (Narayanan et al., 2010).

**Figure 2.**
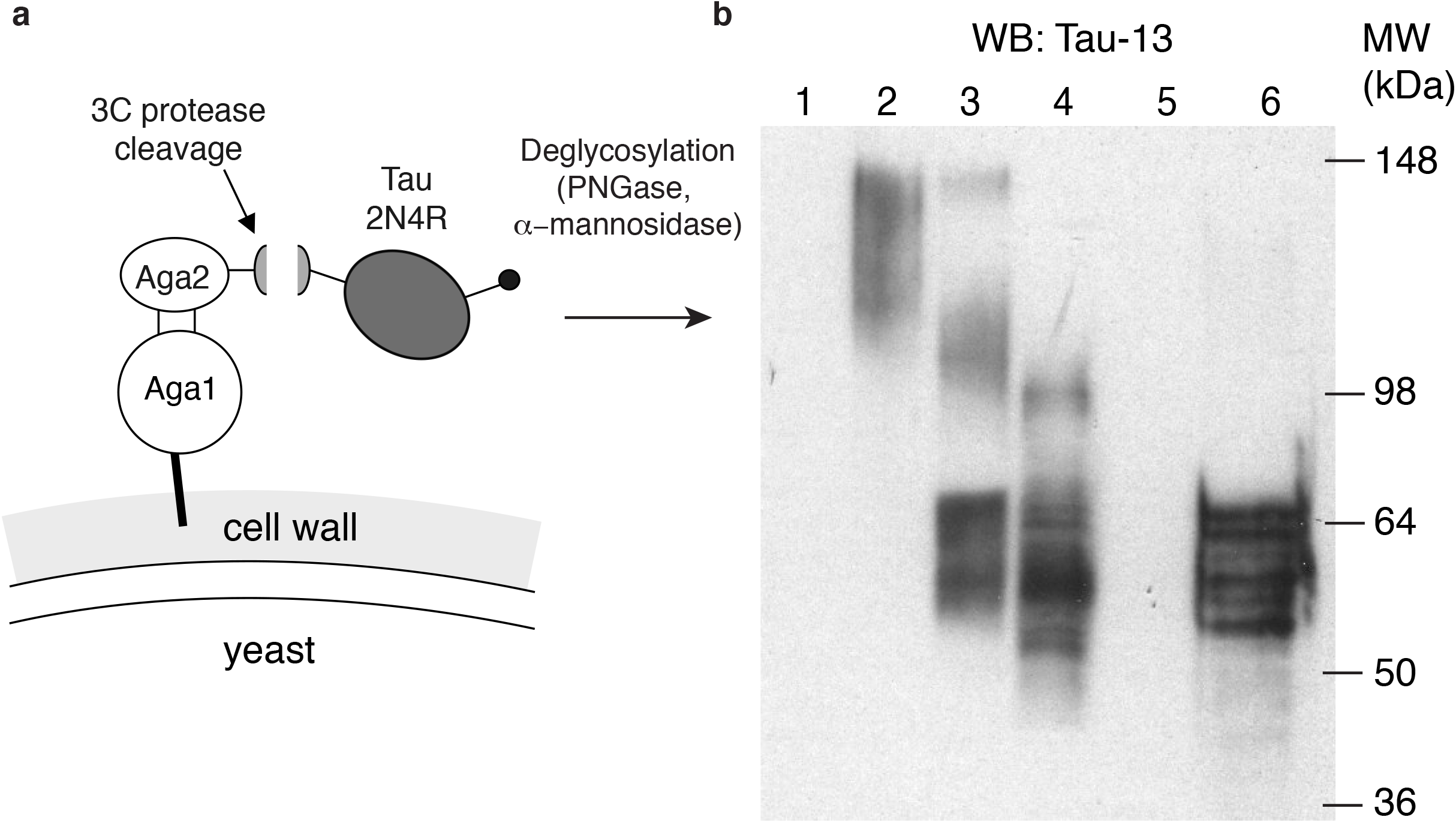
Assessment of glycosylation of yeast-displayed tau using protease cleavage and western blot. (**a**) Schematic representation of yeast surface displayed tau containing a 3C protease cleavage site in between Aga2p and tau. (**b**) Western blot results of 3C protease cleavage products, probed using the anti-tau antibody Tau-13. Samples loaded in lane 1: Yeast cells displaying single-chain variable fragments (scFv) treated with the protease reaction buffer without 3C protease, 2: Yeast cells displaying 2N4R tau treated with the protease reaction buffer without 3C protease, 3: Yeast cells displaying 2N4R tau treated with 3C protease, 4: Yeast cells displaying 2N4R tau treated with PNGase and α1-2,3,6 Mannosidase, followed by 3C protease, 5: Empty, 6: Recombinant tau expressed in *E. coli* (mixture of 6 isoforms).

### Anti-tau antibody binding to yeast-displayed tau

To assess the epitope availability of yeast-displayed 2N4R tau, we assessed its binding to monoclonal anti-tau antibodies with well-defined, phosphorylation-independent linear epitopes (**Fig. 3a**). In addition to the Tau-5 antibody used for initial characterization (**Fig. 1c**), antibodies HT7, Tau-1, and 1E1/A6 showed clear binding to yeast-displayed tau (**Fig. 3b**). However, antibody Tau-46, which binds to the C-terminus of tau, had no binding to yeast-displayed 2N4R tau (**Fig. 3c**, as compared to the control yeast displaying the green fluorescent protein, *P* = 0.268 unpaired t-test), while the c-myc epitope inserted to the C-terminus of tau was clearly detected in the same cells (**Fig. 3c**). To test whether the binding of Tau-46 is impacted by glycosylation of yeast-displayed tau, we treated yeast cells displaying 2N4R tau with PNGase and α-mannosidase. Binding to Tau-46 did not significantly change upon deglycosylation (**Fig. 3c**, compared to non-deglycosylated cells, *P* = 0.157 unpaired t-test), indicating that the absence of binding is not affected by glycosylation of the epitope. These results indicate that in yeast-displayed tau, a wide range of linear epitopes are available for antibody binding, although the C-terminal Tau-46 epitope is unavailable.

**Figure 3.**
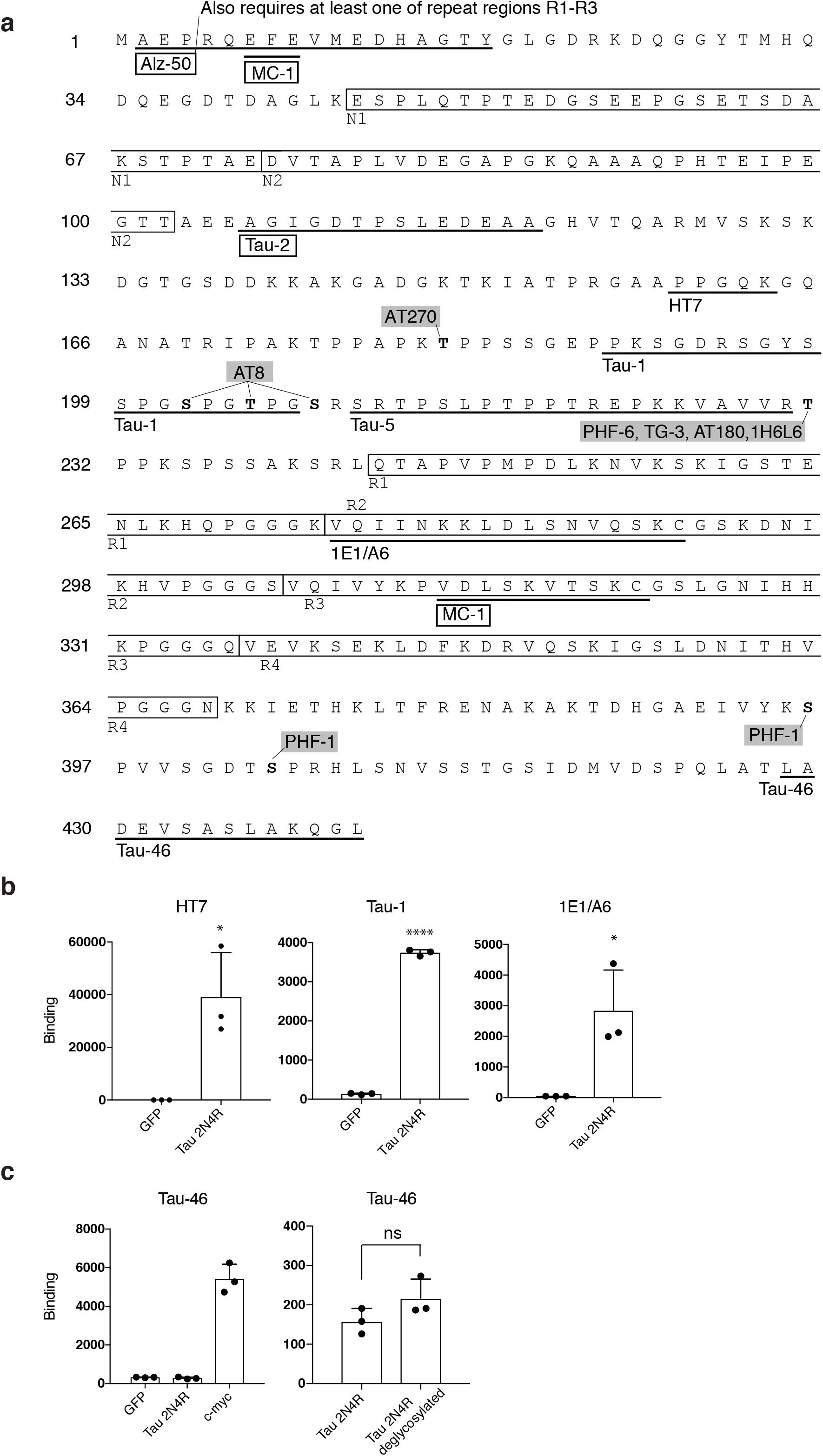
Epitopes of anti-tau antibodies tested against yeast-displayed tau. (**a**) Antibody epitopes mapped on the amino acid sequence of the longest isoform of tau 2N4R. Shaded are antibodies targeting phosphorylation sites, boxes indicate antibodies with discontinuous epitopes. (**b**) Antibody binding to yeast surface displayed GFP or 2N4R tau, quantified using flow cytometry. (**c**) Tau-46 antibody binding to yeast-displayed tau. For each experiment, c-myc labeling was measured to confirm full-length expression of tau. For panels (**b**) and (**c**), data points are geometric means and error bars indicate standard deviation from three independent culture preparations. Statistics for panel (**b**): Unpaired t-test \**P* ≤ 0.05, \*\*\*\**P* ≤ 0.0001. For panel (**c**): Unpaired t-test, n.s. *P* > 0.05 (*P* = 0.157).

### Phospho-tau antibody binding to yeast-displayed tau

Phosphorylation of tau at specific sites is a well-characterized molecular signature for a wide range of neurodegenerative diseases (Hanger et al., 1998; Kopke et al., 1993; Steinhilb et al., 2007). Antibodies specific to phosphorylated tau are widely used to study the mechanism of tau-mediated toxicity and are a promising tool for diagnosis and monitoring disease progression. We assessed the binding of widely used phospho-tau specific antibodies target phosphorylation sites implicated in AD (see **Table 1** for list of antibodies). These antibodies recognize single or multiple phospho-threonine (pT) or phospho-serine (pS) within a specific site in tau. Phospho-tau antibodies AT8 (targeting pS202, pS205, pS208), AT270 (pT181), AT180 (pT231), and PHF-1 (pS396 and pS404) showed no detectable binding to yeast cells displaying 2N4R tau treated with deglycosylating enzymes PNGase and α-mannosidase (**Fig. 4**). In these cells, full-length expression was assessed using the C-terminal c-myc tag (**Fig. 4**). The antibody Tau-1, which binds to non-phosphorylated tau at site P189 – G207 (overlapping with the phospho-sites of AT8) (**Fig. 3**) (Biernat et al., 1992; Carmel et al., 1996), showed clear binding (**Fig. 4**), further indicating absence of phosphorylation. Previous studies showed that human 2N4R tau expressed in the cytosol of yeast *S. cerevisiae* is phosphorylated, and binds to phospho-tau antibodies AT180, AT8, AT270 among others (Vandebroek et al., 2005). The intracellular phosphorylation was driven by the kinase Mds1 (Vandebroek et al., 2005), which is an ortholog of glycogen synthase kinase-3β (GSK-3β) (Puziss et al., 1994). GSK-3β is one of proline-directed protein kinases that phosphorylated tau in neurons (Buee et al., 2000; Lee et al., 2001). The primary reason for the lack of phosphorylation may be the requirement for reducing conditions for these kinases (Puziss et al., 1994). For surface display, since the Aga2p fusion proteins go through the secretory pathway, which is an oxidizing environment (Holkeri & Makarow, 1998; Suh et al., 1999), the yeast-displayed tau is likely to be excluded from the kinase activity.

**Figure 4.**
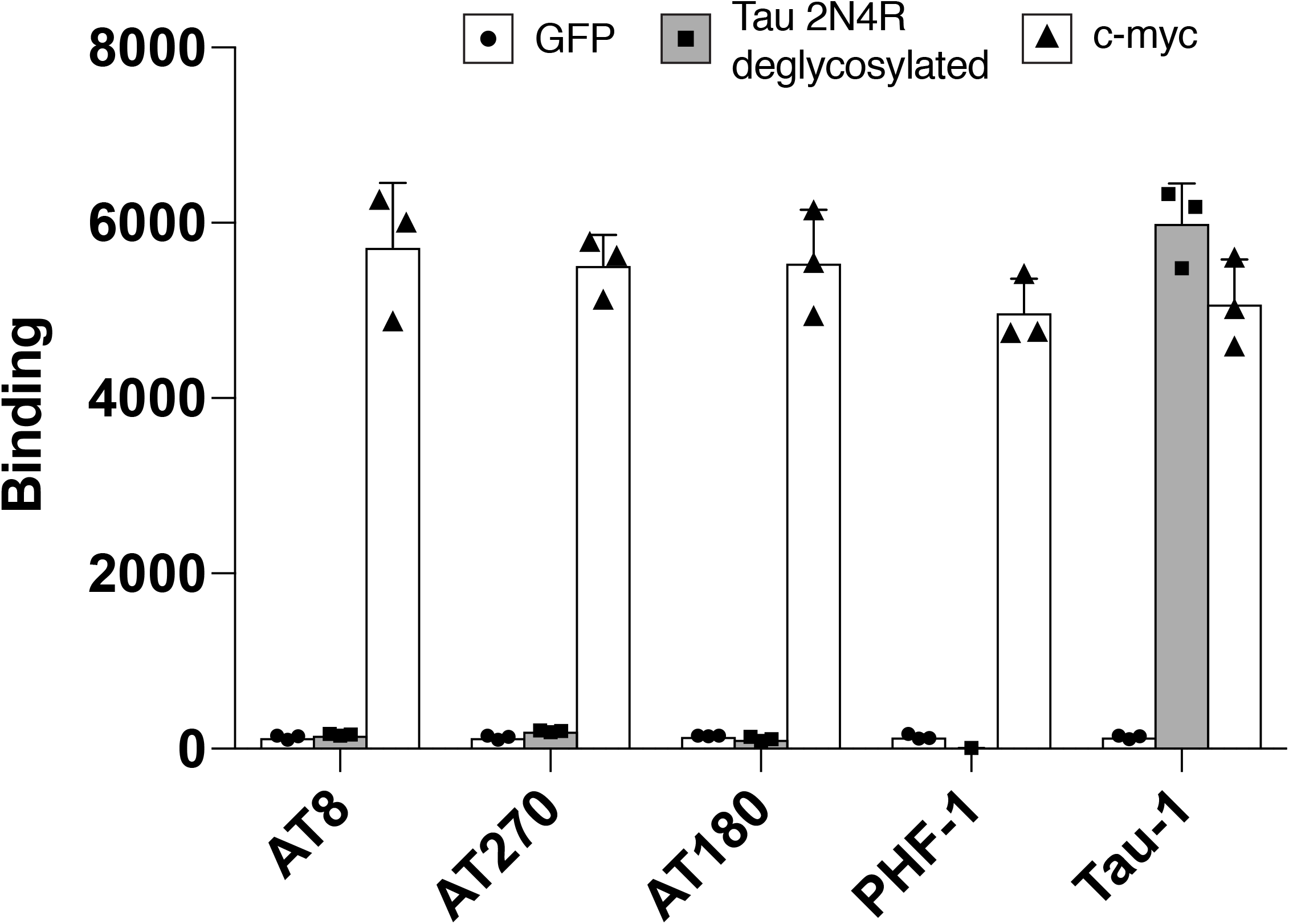
Phospho-tau antibody binding to yeast-displayed tau. Phospho-tau antibody binding to yeast cells displaying either GFP (circles) or deglycosylated 2N4R tau (squares, shaded) was quantified using flow cytometry. Tau-1 has overlapping epitope with AT8, but binds to non-phosphorylated tau. For each experiment, c-myc labeling (triangles) was measured to confirm full-length expression of tau. Data points are geometric means and error bars indicate standard deviation from three independent culture preparations.

### Conformation of yeast-displayed tau

A defining characteristic of tau in patients with Alzheimer’s disease and other tauopathies is the conformational change of tau and formation of paired helical filaments (PHFs) (Iqbal et al., 2009), which are large supramolecular aggregates. Even though tau is mostly unstructured, tau in PHFs possess a structured core and a specific conformation, in which the N-terminus folds in proximity to the repeat domains (Fitzpatrick et al., 2017). Previous results show that tau expressed in the cytosol of *S. cerevisiae* binds to the antibody MC-1 (Vandebroek et al., 2005), which is specific to tau conformation found in PHFs (Carmel et al., 1996; Jicha, Bowser, et al., 1997; Weaver et al., 2000). We assessed the conformation of yeast-displayed tau using conformation-specific antibodies Alz-50 (Wolozin et al., 1986), MC-1 (Jicha, Bowser, et al., 1997), and Tau-2 (Carmel et al., 1996; Watanabe et al., 1992). Previous studies have revealed that Alz-50 and MC-1 have discontinuous epitopes on tau (**Fig. 3a**) (Carmel et al., 1996; Jicha, Bowser, et al., 1997). When we incubated yeast displaying 2N4R tau, all three antibodies showed clear binding (**Fig. 5a**). Furthermore, upon deglycosylation using PNGase and α-mannosidase, the binding to Alz-50 and MC-1 increased by 3- and 5-fold, respectively (**Fig. 5b). These results indicate that the yeast-displayed tau possess conformation found in PHFs. To assess the possibility of aggregation and PHF formation of yeast-displayed tau, we first assessed oligomerization of yeast-displayed tau using tau oligomer-specific antibodies T22 (Lasagna-Reeves et al., 2012) and TOMA-1 (Castillo-Carranza et al., 2014). Yeast cells displaying 2N4R tau did not show significant binding to T22 and TOMA-1, while full-length tau was clearly expressed on the yeast surface (**Fig. 6a****). The absence of binding did not change upon deglycosylation of yeast cells displaying 2N4R tau (**Fig. 6a**), suggesting that the absence of binding to T22 and TOMA-1 was not due to glycosylation. To assess the presence of higher order aggregates of tau, yeast cells displaying 2N4R tau were incubated with Thioflavin S, which stains amyloids including PHF (Yamamoto & Hirano, 1986). Compared to control yeast cells displaying a single-chain antibody variable fragment, no significant difference in staining was detected (**Fig. 6b**). Taken together, these results indicate that yeast-displayed 2N4R tau possesses a conformation characteristic to PHFs, even though it does not form oligomer or aggregates. It is known that tau in PHF conformation can be monomeric (Weaver et al., 2000) and changes conformation prior to aggregation (Carmel et al., 1996). Unlike tau expressed in the yeast cytosol, which was hyperphosphorylated and showed MC-1 binding (Vandebroek et al., 2005), we did not detect any phosphorylation at major phosphorylation sites (**Fig. 4**). Hyperphosphorylation causes conformational change of tau (Augustinack et al., 2002; Hanger et al., 2009), but our results suggest a different mechanism for conformational change in yeast-displayed tau. One possible driving force may be the high density of anionic surface charge in yeast cell surface. Yeast plasma membrane consists of negatively charged phospho-lipid head-groups (van der Rest et al., 1995), and the cell wall is also negatively charged due to phosphorylation of the mannosyl side chains (Lipke & Ovalle, 1998). Previous studies show that purified non-phosphorylated tau forms PHFs upon incubation with sulfated glycosaminoglycans, such as heparin (Goedert et al., 1996). Subsequent studies have shown conformational change of recombinant tau driven by polyanions, such as sulfate containing polymers with a various sugar backbone (Hasegawa et al., 1997), polyglutamate (Friedhoff et al., 1998), nucleic acids such as RNA (Hasegawa et al., 1997; Kampers et al., 1996), and fatty acid micelles (Wilson & Binder, 1997). In general, tau conformational change can be driven by increasing the ionic strength (Crowther et al., 1994; Wille et al., 1992). Thus, due to the inherent negative surface charge, the yeast surface display approach provides a unique platform for characterizing protein interactions to pathogenic conformation of human tau.

**Figure 5.**
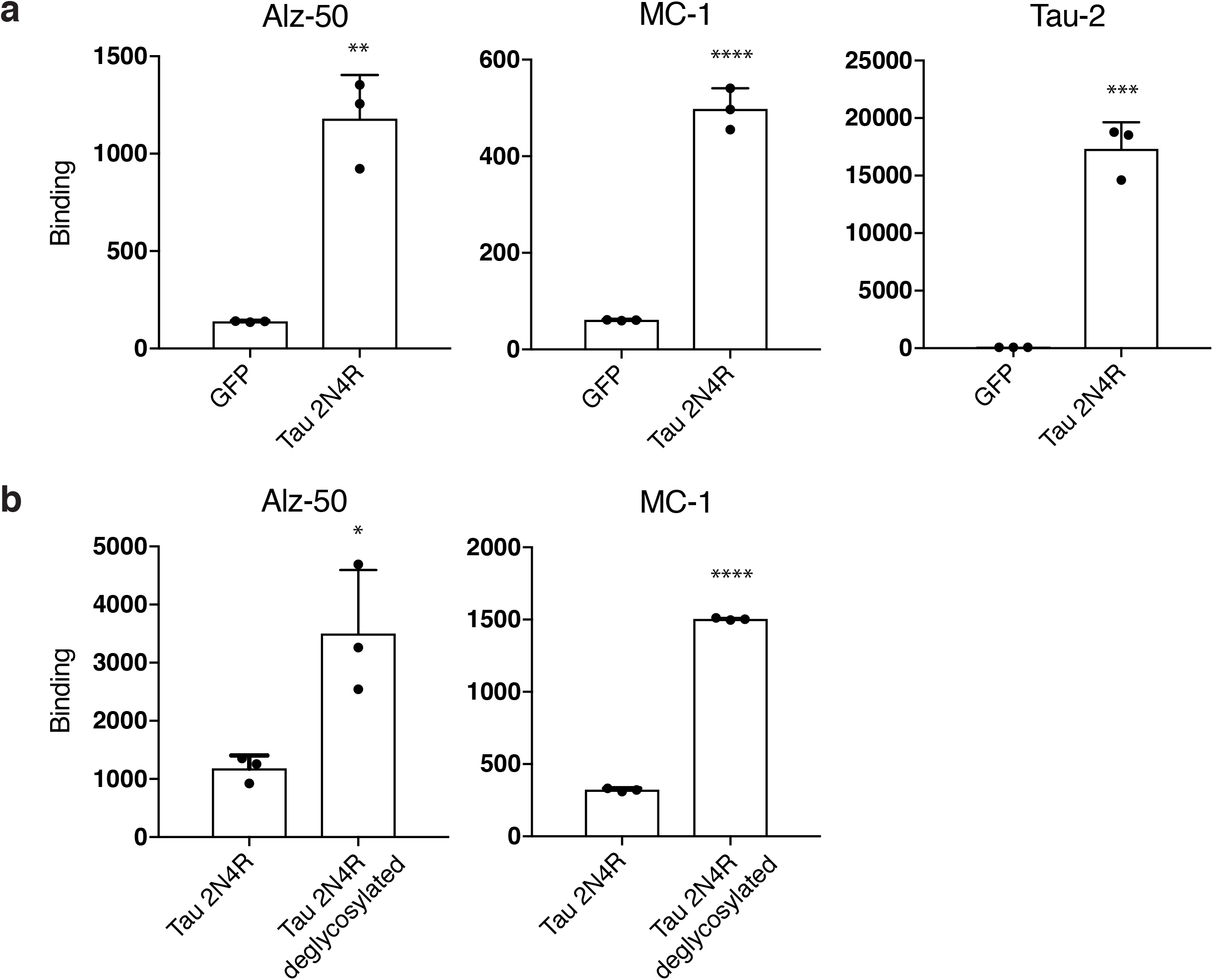
Yeast surface displayed tau binds to antibodies specific to conformation found in Alzheimer’s disease patients. (**a**) Antibody binding to yeast surface displayed GFP or 2N4R tau, quantified using flow cytometry. Alz-50, MC-1 and Tau-2 are conformation-specific antibodies. (**b**) Enhancement of Alz-50 and MC-1 antibody binding upon deglycosylation of yeast-displayed 2N4R tau. Data points are geometric means and error bars indicate standard deviation from three independent culture preparations. Statistics: Unpaired t-test \**P* ≤ 0.05, \*\**P* ≤ 0.01, \*\*\**P* ≤ 0.001, \*\*\*\**P* ≤ 0.0001.

**Figure 6.**
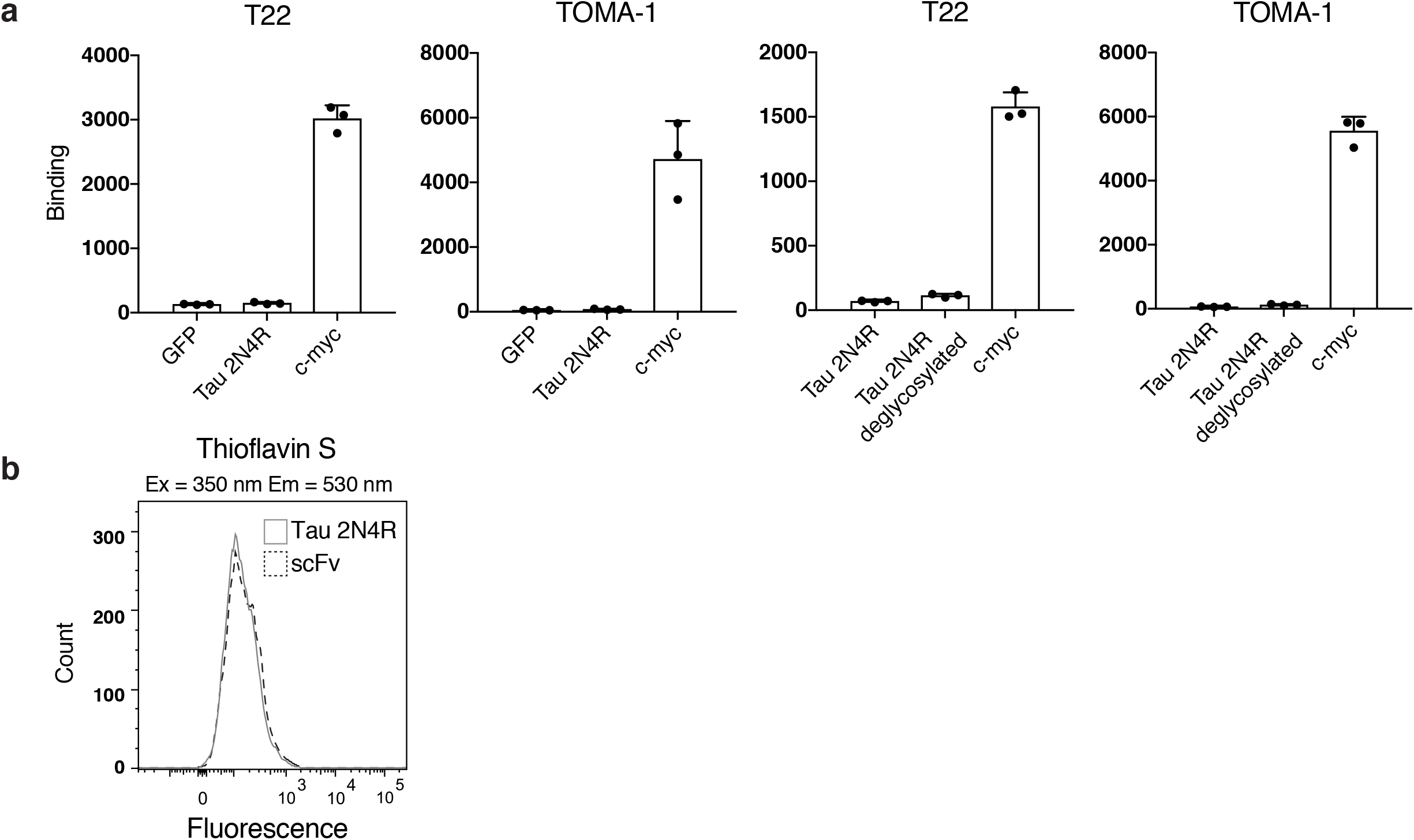
Assessment of oligomer and aggregate formation of yeast surface displayed tau. (**a**) Antibody binding to yeast surface displayed GFP, 2N4R tau, or deglycosylated 2N4R tau quantified using flow cytometry. Antibodies T22 and TOMA-1 are specific to tau oligomers. For each experiment, c-myc labeling was measured to confirm full-length expression of tau. Data points are geometric means and error bars indicate standard deviation from three independent culture preparations. (**b**) Thioflavin S staining of yeast cells displaying either a single-chain antibody variable fragment (scFv) or 2N4R tau. Green fluorescence (530 nm emission) of Thioflavin S under ultraviolet stimulation (350 nm excitation), which increases upon paired helical filament aggregate binding, was measured using flow cytometry.

## Conclusions

We have shown that full-length human tau, including the longest isoform found in the adult brain (2N4R) was displayed on the surface of yeast *S. cerevisiae*. Yeast-displayed tau was glycosylated, which was removed using PNGase and α-mannosidase. Many linear epitopes in tau, ranging from the N-terminus to the 4R repeat region were available for antibody binding, except the C-terminal epitope targeted by the antibody Tau-46. Unlike human tau expressed in the cytosol of *S. cerevisiae*, yeast-displayed tau did not show any phosphorylation in phospho-sites important for conformational change (those targeted by antibodies AT8, AT270, AT180, PHF-1, and Tau-1). However, yeast-displayed tau showed conformation recognized by antibodies specific to PHF conformation, Alz-50, MC-1, and Tau-2, potentially due to the negative charge on the yeast surface. Even though the tau possessed a conformation found in PHFs, oligomerization or aggregation into larger filaments were undetected. We anticipate that the yeast-displayed tau will be a valuable tool for studying its interaction with other proteins, and potentially characterizing its pathogenic conformation.

## Acknowledgements

This work was funded by the NSF grant 1706743. The authors have no conflict of interest.

